# Resting-state based prediction of task-related activation in patients with disorders of consciousness

**DOI:** 10.1101/2021.03.27.436534

**Authors:** Michael M Craig, Ioannis Pappas, Judith Allanson, Paola Finoia, Guy Williams, John D Pickard, David K Menon, Emmanuel A Stamatakis

## Abstract

**Background:** Assessment of the level of awareness of people with disorders of consciousness (DOC) is clinically challenging, motivating several studies to combine brain imaging with machine learning to improve this process. While this work has shown promise, it has limited clinical utility, as misdiagnosis of DOC patients is relatively high. As machine learning algorithms rely on accurately labelled data, any error in diagnosis will be learned by the algorithm, resulting in an equally limited diagnostic tool. The goal of the present study is to overcome this problem by stratifying patients, not by diagnosis, but by their capacity to perform volitional tasks during functional magnetic resonance imaging (fMRI) scanning.

**Methods:** A total of 71 patients were assessed for inclusion. They were excluded for the final analysis if they had large focal brain damage, excessive head motion during scanning, or suboptimal MRI preprocessing. Patients underwent both resting-state and task-based fMRI scanning. Univariate fMRI analysis was performed to determine if an individual patient had brain activity consistent with having retained volitional capacity (VC). Differences in resting brain network connectivity between patients with VC and patients without volitional capacity (non-VC) were measured. Connectivity data was then entered as input to a deep learning framework. We used a deep graph convolutional neural network (DGCNN) on connectivity data to identify a specific brain network that most significantly differentiates patients.

**Findings:** We included 30 patients in our final analysis. Univariate analysis revealed that 13 patients displayed signs of VC, while 17 did not. We found that resting-state connectivity between frontoparietal control and salience network was significantly different between VC and non-VC patients (T(28) = 3.347, p = 0.0023, Bonferroni corrected p = 0.042). Furthermore, we found that using frontoparietal control network connectivity as input to the DGCNN resulted in the best classification performance (test accuracy = 0.85; ROC AUC = 0.92).

**Interpretation:** We found that the DGCNN performed best at discriminating between patients with VC when using only the frontoparietal control network as input to the model. The use of this deep learning method is a significant advance since its inherent flexibility permits the inclusion of both whole-brain and network-specific properties as input, allowing us to classify patients as either having or not having VC. This inclusion of multi-scale inputs (e.g. whole-brain and network-level) facilitates model interpretability and increases our understanding of the neurobiology of DOC. The results propose that the integrity of frontoparietal control network, a brain network well known to play a key role in executive functions and cognitive control, is essential for volitional capacity preservation in patients with DOC. The study also lays groundwork for development of a biomarker to aid in the diagnosis of DOC patients.

**RESEARCH IN CONTEXT:** *Evidence before this study:* Disorders of consciousness (DOC) are a group of severe brain disorders characterised by damage to the neural systems underlying wakefulness and awareness. DOC are often caused by traumatic brain injury, hypoxia, or neurodegenerative diseases. The motor and cognitive impairments in DOC patients make providing an accurate diagnosis very challenging. Diagnosis is primarily made at the bedside by assessing a patient’s response to motor commands.

## INTRODUCTION

Disorders of consciousness (DOC) are a severe group of conditions characterised by perturbations to the neural systems underlying awareness and wakefulness (Giacino, Fins, Laureys, & Schiff, 2014). These clinical states are often the result of relatively focal traumatic brain injury or more global injury caused by hypoxia. In coma, patients lie with their eyes closed and do not respond to any external stimulation. Patients whose eyes are open, but are still unable to respond to external stimuli are said to have unresponsive wakefulness syndrome (UWS), also known as vegetative state (Jennet & Plum, 1972; Laureys et al., 2010). When patients are capable of producing transient, but reproducible non-reflexive behaviours they are said to be in minimally conscious state (MCS; Giacino et al., 2002). Assessment of awareness levels in these patients is critical not only for guiding treatment and refining prognostication, but also for providing reliable information to their families. Diagnosis is primarily made at the bedside by observing a patient’s motor responses to a variety of commands with standardised tests such as the Coma Recovery Scale-Revised (CRS-R; Giacino et al., 2014; Giacino, Kalmar, & Whyte, 2004). However, accurately determining the level of awareness in these patients is complicated since they often suffer from additional cognitive or sensory impairments. These challenges have motivated the development of neuroimaging biomarkers that can aid in the diagnosis and prognosis of patients with DOC. Moreover, with the increased interest in applying deep learning-based machine learning models to healthcare, it is critical to explore how these methods can be used to improve DOC patient care and deepen our understanding of neuropathology associated with this disorder.

To this end, Demertzi et al. (2015) combined resting-state functional connectivity with a support vector machine (SVM) classifier to classify patients with different levels of DOC. This study demonstrated that functional connectivity measures have significant discriminatory power when assessing DOC patients (Demertzi et al., 2015).

Studies utilising MRI to assess DOC patients either measure activity or connectivity during resting-state (Boly et al., 2009; Demertzi et al., 2014; Di Perri et al., 2016), sensory stimulation (Boly et al., 2007; Schiff et al., 2005) or during active mental imagery paradigms (Bardin, Schiff, & Voss, 2012; Menon et al., 1998; Monti et al., 2010; Owen et al., 2006). Mental imagery paradigms are of particular interest, as this work has shown that a subset of DOC patients can voluntarily respond to commands while in the scanner. A seminal study by Owen et al. (2006) identified a patient who fulfilled all the clinical criteria for UWS but was able to respond to commands by modulating her fMRI activity during a behavioural task. While these approaches demonstrate the utility of using mental imagery tasks to assess volitional ability in patients, they are limited by the fact that many institutions lack the facilities and technical expertise to implement these paradigms and the associated analyses. Additionally, several issues are introduced when using task-based fMRI in a clinical setting, including practice effects and habituation that can result in weaker signal upon repeat exposure to the same task (Greicius et al., 2008). More importantly, though, DOC patients often suffer from irregular circadian rhythms and fluctuating arousal, which can correlate with increased or decreased attention and cognitive ability (Blume et al., 2017). To this end, recent evidence has shown that CRS-R scores taken from the same patients were higher in the morning than in the evening (Cortese et al., 2015). This same problem would likely affect performance in an fMRI-based task. This makes using resting-state scanning, which does not rely on task engagement/performance, a better option for identifying a biomarker.

Taken together, a useful clinical tool for extending our understanding of DOC patients would provide information on the patient’s internal state using a simple measurement like resting-state fMRI. Ideally, this tool would not be a black box but would have a degree of interpretability based in cognitive neuroscience. For this reason, the present study utilises resting-state functional connectivity and deep learning, not to provide a specific diagnosis, but to classify patients with DOC as being able or unable to respond to volitional mental imagery tasks. This decision is motivated by the fact that a clinical diagnosis will always be accompanied with extensive behavioural evaluation, and therefore stratifying patients by volitional ability provides clinicians with more actionable information to aid diagnosis, and not simply reinforce their current assessment. Resting-state fMRI data were collected from patients who are able (VC group) or unable (non-VC group) to respond to a volitional mental imagery task. We then applied functional connectivity analyses to determine if there were significant differences in network dynamics between VC and non-VC patients, providing us with some grounding in basic cognitive network neuroscience. Finally, we implemented a cutting-edge deep graph convolutional neural network (DGCNN) to automatically differentiate between groups and to determine whether a specific brain network provided the most discriminative capacity between patients capable of performing the volitional mental imagery task.

## METHODS

### Patients

A sample of 71 DOC patients were considered for inclusion in this study. Patients were recruited from specialised long-term care centres. To be invited to the study, patients must have had a DOC diagnosis for at least 3 months, consent from their legal representative, and were clinically stable and appropriate for ambulance transfer to Addenbrooke’s Hospital. The exclusion criteria included any medical condition that made it unsafe for the patient to participate (decision made by clinical personnel blinded to the specific aims of the study or any reason they were unsuitable to enter the MRI scanner environment (e.g. non-MRI-safe implants)). After admission, each patient underwent clinical and neuroimaging testing. Patients spent a total of 5 days (including arrival and departure days) at Addenbrooke’s Hospital. Patients were systematically excluded from the final cohort based on the following: 1) large focal brain damage (i.e. more than 1/3 of one hemisphere); 2) excessive head motion during resting-state scanning (i.e. greater than 3mm in translation and/or 3 degrees in rotation); 3) suboptimal segmentation and spatial normalization of images. This process left us with a total of 30 patients (Figure 1A; VS = 13, MCS = 17). The VC group had 3 VS patients and 10 MCS patients, two of whom were diagnosed as eMCS. The non-VC group had 9 VS patients and 8 MCS patients, none of whom were eMCS (A complete overview of patient information can be found in Table 1). Patients had an age range of 19-70 (mean = 38.2, std. dev. = 15.8; mean VC = 37.5, std. dev. = 15.6; mean non-VC = 38.8, std. dev. = 16.5).

**Figure 1:**
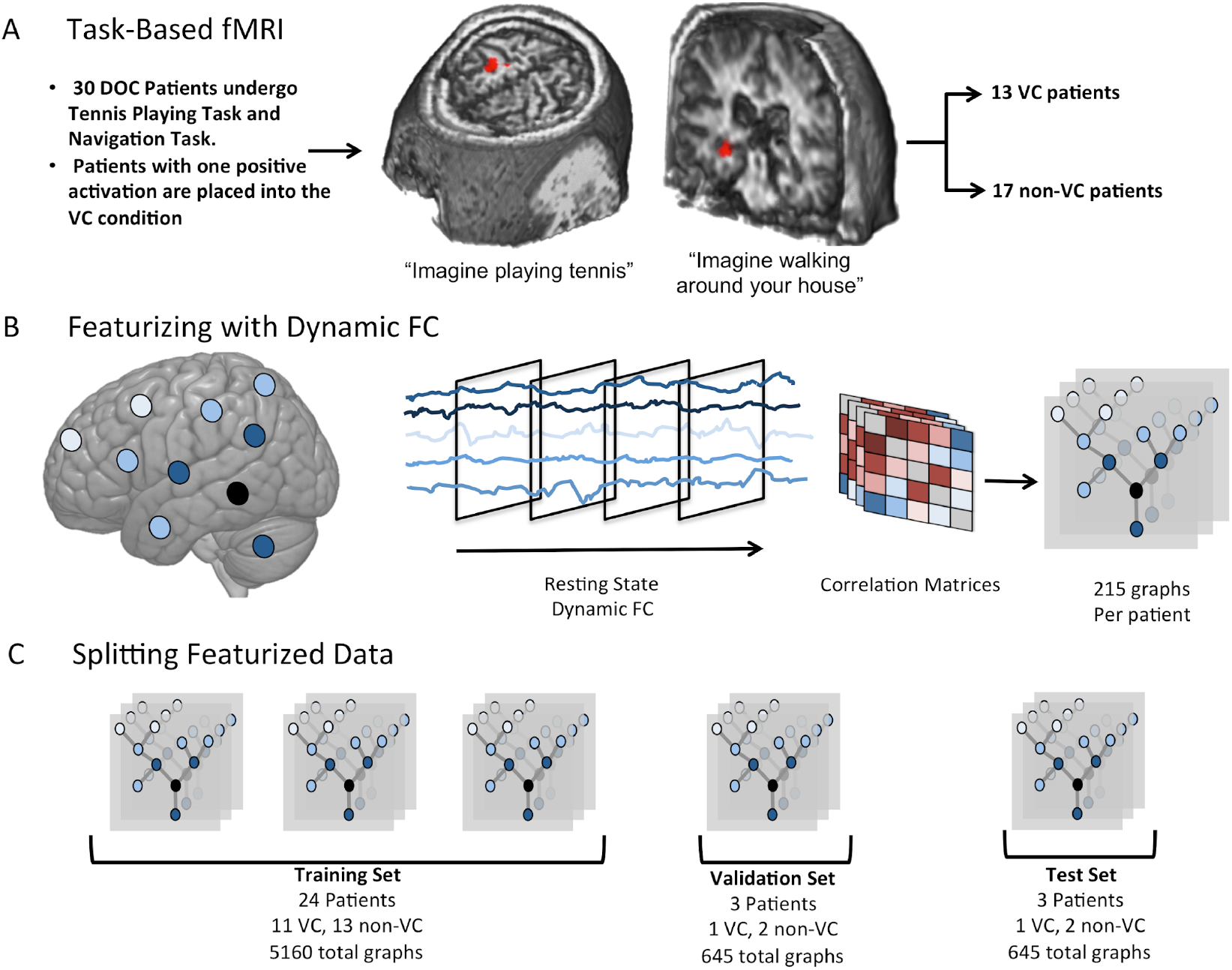
Study Overview. This figure provides an overview of several key elements of the experimental design. **A** shows that task-based fMRI using the previously described mental imagery paradigms. This analysis was used to separate patients with VC and non-VC patients into separate conditions for our classification model. **B** shows how we used dynamic functional connectivity to augment our brain network data. This allowed us to generate 215 datapoints per patient, resulting in a dataset large enough to train a deep learning model. **C** shows how we split our data into training, validation and test sets. Importantly, this split was done per-patient, as mixing each patient’s data into the three sets would result in over-fitting.

**Table 1:** Clinical information table on patients who were selected to be in the final analysis. (MCS = Minimally Conscious State; VS = Vegetative State; SD = Severely Disabeled)

| ID | Team Dx | Age at scan | CRS | Response | Aetiology |
| --- | --- | --- | --- | --- | --- |
| 1 | SD | 22 | 20 | Yes | TBI |
| 2 | VS | 46 | 6 | No | TBI |
| 3 | MCS- | 57 | 12 | No | TBI |
| 4 | EMCS | 47 | 14 | Yes | TBI |
| 5 | EMCS | 61 | 16 | Yes | TBI |
| 6 | MCS | 54 | 10 | Yes | TBI |
| 7 | VS | 19 | 8 | Yes | Anoxic |
| 8 | MCS- | 31 | 10 | No | Anoxic |
| 9 | MCS | 38 | 11 | Yes | TBI |
| 10 | VS | 24 | 8 | No | TBI |
| 11 | MCS | 29 | 10 | No | TBI |
| 12 | MCS | 23 | 7 | Yes | TBI |
| 13 | MCS | 19 | 12 | No | TBI |
| 14 | MCS | 70 | 9 | No | Cerebral bleed |
| 15 | VS | 57 | 7 | No | TBI |
| 16 | MCS- | 30 | 9 | Yes | Anoxic |
| 17 | VS | 36 | 8 | Yes | Anoxic |
| 18 | VS | 39 | 8 | Yes | TBI |
| 19 | VS | 23 | 7 | No | Anoxic |
| 20 | MCS+ | 24 | 13 | Yes | TBI |
| 21 | VS | 40 | 7 | No | Anoxic |
| 22 | VS | 62 | 7 | No | Anoxic |
| 23 | VS | 46 | 5 | No | Anoxic |
| 24 | MCS | 21 | 11 | No | TBI |
| 25 | MCS- | 67 | 11 | Yes | TBI |
| 26 | VS | 55 | 12 | No | Hypoxia |
| 27 | MCS | 30 | 8 | No | TBI |
| 28 | MCS | 28 | 8 | Yes | TBI |
| 29 | MCS | 22 | 10 | No | TBI |
| 30 | VS | 28 | 6 | No | ADEM |

On the day of scanning, patient’s behaviours were observed and recorded using the CRS-R (Giacino et al., 2004). This is a commonly used measure of awareness in DOC patients (Seel et al., 2010). This measure is split into 23 items over 6 domains to assess behaviours of DOC patients. The test was administered by a trained neuropsychologist or medic. Factors such as the time of day and patient postural position were also noted.

### Mental Imagery Tasks

The mental imagery tasks used here have been previously used to assess awareness in DOC patients (Owen et al., 2006; Monti et al., 2010; Bardin et al., 2012) and have been validated in healthy participants (Fernandez-Espejo et al. 2014). Briefly, each patient was asked to perform two mental imagery tasks. In the motor imagery task, the patients were instructed to imagine being on a tennis court, swinging their arm to hit the ball back and forth with an imagined opponent. The spatial imagery task involved the patient to imagine walking to each room of their home or navigating the streets in a familiar city, and to picture what they would observe if they were there. Each task consisted of five rest-imagery block cycles (30 seconds each). Each rest block was cued with the spoken word “relax” and imagery block was cued with the spoken word “tennis” or “navigation”.

### Data Acquisition

Data was acquired at Addenbrooke’s Hospital in Cambridge, UK, on a 3T Tim Trio Siemens system (Erlangen Germany). T1-weighted images were acquired with an MP-RAGE sequence (TR = 2300ms, TE = 2.47ms, 150 slices, 1 x 1 x 1.2mm^2^ resolution). Functional images were acquired using an echo planar sequence (TR = 2000ms, TE = 30ms, flip angle = 78°, 32 slices, 3 x 3 x 3.75mm^2^ resolution).

### Data Preprocessing and Analysis

#### Brain Network Parcellation

To model brain networks as graphs for input into our graph-based deep learning model, we first parcellated the cortex into different brain regions. We used the 129-region Lausanne Parcellation (Hagmann et al., 2008) which consists of 64 regions per hemisphere and 1 bilateral brain stem region. A central aim of our study was to identify which large scale brain networks are important for differentiating between patient groups so we assigned each brain region to a large scale brain network using network masks (Smith et al., 2009). To assign each ROI to one of the networks, we calculated the overlap of each region with the mask of each network. The maximum overlap was used to assign the ROI to the respective network. Previous work has shown that interactions between high-level cognitive networks are important in stratifying DOC patients (Di Perri et al., 2016) especially the DMN, FPCN and SN. These networks resulted in a total of 71 ROIs included in our final analysis.

### High Resolution T1 and functional MRI Data Preprocessing

Preprocessing was performed with Statistical Parametric Mapping 12 (SPM12; http://www.fil.ion.ucl.ac.uk/spm/) and MATLAB version 2017a (http://www.mathworks.co.uk/products/matlab/). The first five volumes for each sedation condition were removed to eliminate saturation effects and achieve steady-state magnetization. The fMRI images were then subjected to slice timing correction and realignment to correct for movement. Participant’s high-resolution structural images were coregistered to the mean EPI (produced from the realignment process) and segmented into grey matter, white matter and cerebrospinal fluid masks (Ashburner & Friston, 2005). Next, the images were normalised and resampled to Montreal Neurological Institute (MNI) space with a resolution of 2 x 2 x 2 mm^3^. Functional images were smoothed with a 6mm FWHM Gaussian kernel. To further reduce movement-related and physiological artefacts, data underwent de-spiking with a hyperbolic tangent squashing function. Next, the aCompCor technique was used to remove the first 5 principal components of the signal from the white matter and cerebrospinal fluid masks, as well as 6 motion parameters and their first order temporal derivatives and a linear de-trending term (Behzadi, Restom, Liau, & Liu, 2007). Functional images were highpass filtered to remove very low-frequency fluctuations associated with scanner noise (0.009 Hz < *f*) (Craig, Manktelow, Sahakian, Menon, & Stamatakis, 2018).

#### Univariate fMRI Analysis

Univariate fMRI analysis was conducted on all 30 patients for both the motor and spatial mental imagery tasks. The analyses were performed using FSL (version 5.0.9). For each functional scan a general linear model consisting of contrasting periods of rest and active imagery was computed. Results were considered significant at a cluster level of z > 2.3 (corrected p < 0.05) (Monti et al. 2010).

#### Functional Connectivity Analysis

Functional connectivity was calculated by measuring the Pearson correlation coefficient between each of the 71 regions in DMN, FPCN and Salience networks. This allowed us to examine both within-network and between-network connectivity. These measures were calculated by averaging the correlation values within or between each network of interest. We did not threshold these network measures to preserve any anticorrelation between networks of interest. An independent-sample t-test was used to test for significant differences between the positive and negative volitional responder groups in the DOC dataset.

### Development of Deep Learning Framework

For input into our model, we split the dataset into a training (n=24), validation (n=3) and test set (n = 3). The validation dataset was necessary to tune the model’s hyperparameters which optimises performance. Because deep learning methods need a large amount of training data, we augmented our data using dynamic functional connectivity. Using a sliding window approach, we calculated one graph for every 80 volumes (Figure 1B). With each resting-state scan consisting of 300 functional volumes (with the first 5 volumes removed), dynamic functional connectivity resulted in 215 graphs per patient (training set = 5160; validation set = 645; test set = 645; Figure 1C).

The DGCNN is adapted from (Zhang, Cui, Neumann, & Chen, 2018; https://github.com/muhanzhang/DGCNN) and consists of three sequential stages. 1) Graph convolutional layers to extract node connectivity features; 2) a SortPooling layer to sort node features and equate input features size for, 3) a series of classical convolutional and fully connected neural network layers to read the sorted graph representations and make predictions (Zhang et al., 2018).

The graph convolutional layers aggregate node information from neighbouring nodes to extract multi-scale graph substructures important for classification (See Figure 2 for schematic overview). For each graph convolutional layer, given a graph **A** (a binary functional connectivity matrix) and its node information, information is propagated through the network with:

**Figure 2:**
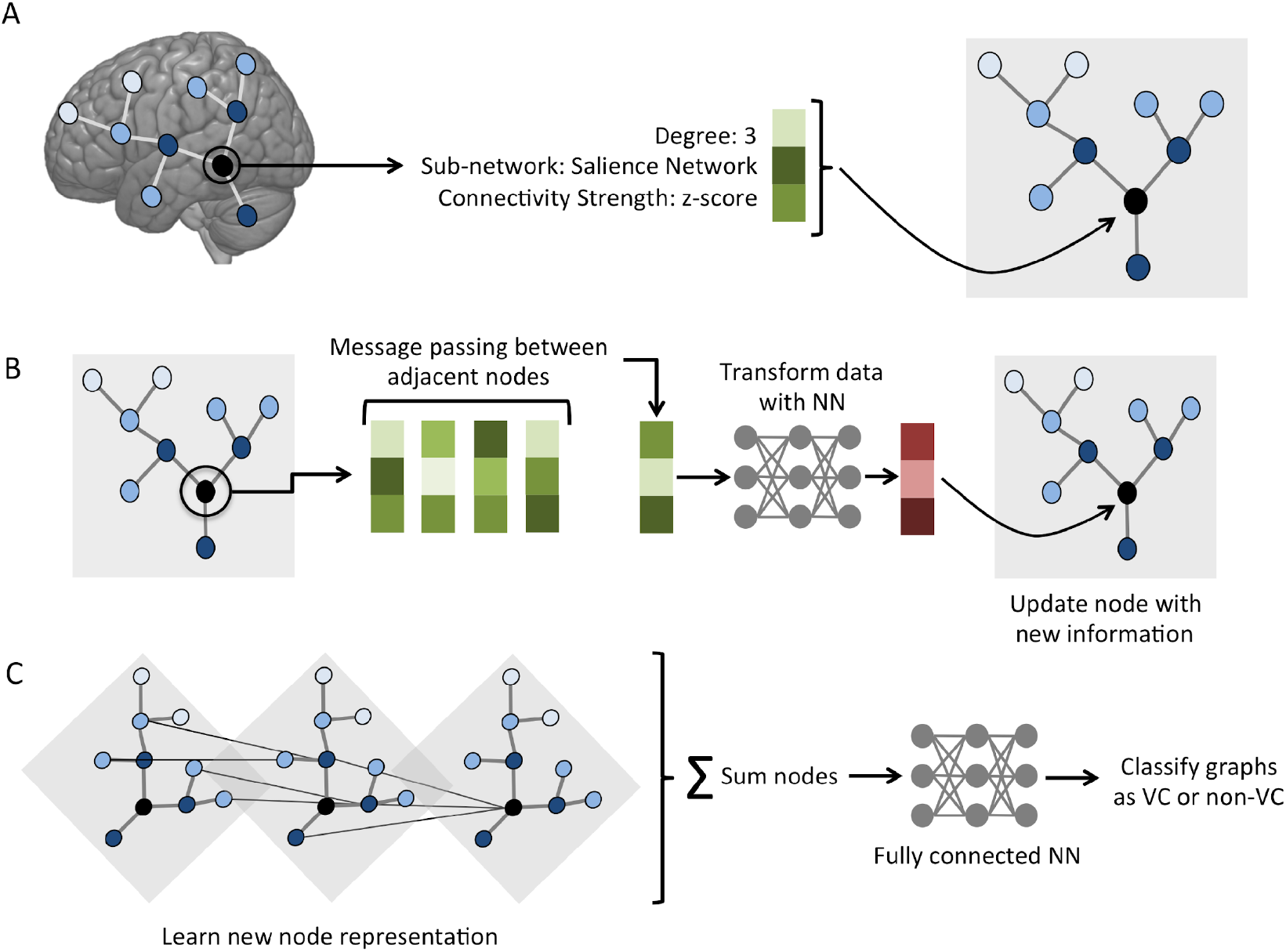
Schematic of graph convolutional architecture. **A** shows that each node in the graph is represented by a feature vector. The feature vector contains information about the node, including degree, which sub-network it belongs to, and the summed connectivity strength with adjacent nodes. **B** demonstrates how each node iteratively collects information (passes messages) about adjacent nodes, which is then transformed by a neural network into new information at the initial node. **C** shows that each node can collect multi-scale information by stacking graph convolutional layers. At the final layer, information is summed and passed into a fully connected neural network to classify the graph as belonging to either the VC or non-VC class.

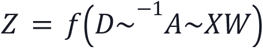

where *A*∼ =*A* + *I* is the adjacency matrix plus the identity matrix. Inclusion of the identity matrix results in self-loops, meaning that for the node undergoing convolution, it is multiplying its neighbours and its own feature vectors 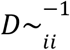 is the inverse of the diagonal degree matrix. We take the inverse of this matrix to normalise *A*∼ therwise *A*∼ will change the scale of the feature vectors *W ∈ R^cxc’^* is a matrix of trainable weights and *f* is a nonlinear activation function.

The convolution itself contains four separate steps. The first step is a linear feature transformation applied to the node feature matrix by **XW**. This maps the *c* feature channels to the *c’* feature channels in the next layer and the trainable weights are shared amongst all nodes. The next step, *A*∼*Y*, where *Y*:=*XW*, passes the neighbouring node’s feature vectors (plus the self-loop) to the node that is undergoing convolution. Step three is the normalisation of row *i* by multiplication with 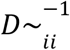. The final step applies a nonlinear activation function *f* (in this case a rectified linear unit or ReLU which outputs the final graph convolution result.

The preceding four steps aggregate *local* nodal information. To extract multiscale substructures from the data, thus resulting in a richer featurisation and ultimately better classification, we stack multiple graph convolutional layers on top of one another. This is done using:

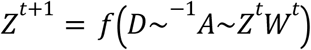

where 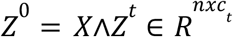 is the output of the *t*^th^ graph convolutional layer. Here, *c*_*t*_ is the number of output channels at layer *t* and the weight matrix 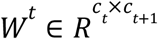 maps channels in the current layer to *c*_*t+1*_ channels in the following layer. The final concatenated output represents a multi-scale feature descriptor for each node in the graph.

The SortPooling layer sorts the output of the graph convolutional layer so that node feature vectors are pooled together and outputted in a consistent order. This is important because the final 1D convolutional step is most effective at classification when features are presented in a consistent order (Zhang et al., 2018). This final step consists of two layers of classical 1D convolutional layers, each with a convolutional layer, followed by a rectified linear unit activation function and maxpooling layer. This is followed by a fully connected layer and a softmax layer for classification.

We used Bayesian hyperparameter optimisation in python using pyGPGO (http://pygpgo.readthedocs.io/en/latest/) as opposed to grid search to optimise our models. Briefly, Bayesian optimisation uses previous results from the model to compute the posterior expectation of the loss function and then uses this expectation to find the next optimal set of hyperparameters to improve the model (Snoek et al., 2015). We performed Bayesian optimisation by training our model many times with increasingly optimised hyperparameters and evaluating its accuracy on the validation dataset. The best set of hyperparameters for each model was then used to make predictions on the final held-out test dataset.

To assess the results of each binary classification we used the following metrics: Precision (True positives/True Positives + False Positives), recall (True Positives/True Positives + False Negatives), F1 Score (Harmonic mean of Precision and Recall), and the area under the receiver operating characteristic curve (ROC-AUC; Bishop, 2006). The ROC curve is a plot of True Positive Rate against False Positive Rate for different cut-offs of a diagnostic test and is a measure of the trade-off between sensitivity (true positive rate) and specificity (1 – false positive rate). As our analysis has balanced classes (i.e. an equal number of examples for each cognitive state), the ROC-AUC was considered the most important metric (Davis & Goadrich, 2006).

## RESULTS

### Positive and Negative Responders to Mental Imagery Tasks

We found a total of 13 patients who had a positive response to at least one mental imagery task (**Figure 3**). Patient 20 responded positively to both the motor imagery and spatial imagery tasks. 10 patients had positive responses to the spatial imagery task, while 4 patients had positive responses to the motor imagery task. This resulted in having 13 patients in the VC group and 17 patients in the non-VC group. We also found that CRS-R scores were significantly greater in responsive (median = 10.0, Interquartile Range = 5.0) versus unresponsive (median = 8.0, Interquartile Range = 3.0) patients (Mann-Whitney U = 68.0, p = 0.0381).

**Figure 3:**
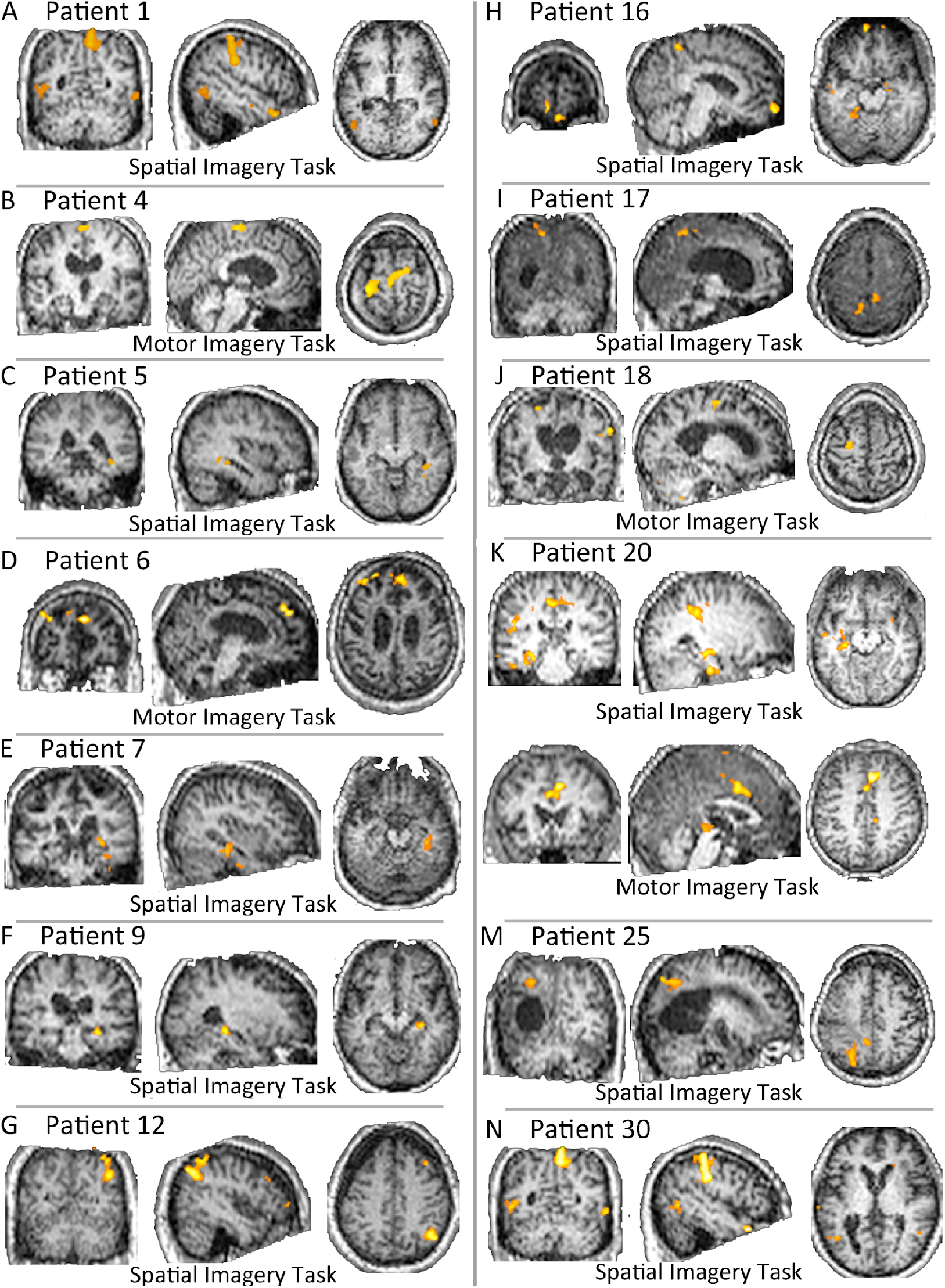
Activity of VC patients. We found a total of 13 patients who responded to one or more mental imagery tasks. Patient 20 had positive responses to both the spatial and motor imagery task.

### DMN, FPCN and Salience Network Functional Connectivity in Disorders of Consciousness

Previous work has shown that resting-state functional connectivity between DMN and putative task-positive networks is altered in patients with disorders of consciousness (Di Perri et al., 2016). We sought to expand on this work by examining within and between DMN connectivity, as well as within and between FPCN and Salience network connectivity in patients with DOC.

We did not observe any significant differences for within DMN connectivity (T(28) = 0.203, p = 0.84; **Figure 4A**), FPCN connectivity (T(28) = −1.681, p = 0.104; **Figure 4B**), or Salience network connectivity (T(28) = 0.732, p = 0.103; **Figure 4C**) for patients who responded to volitional tasks compared to those who did not. We did not observe any significant differences in DMN-FPCN connectivity (T(28) = 0.482, p = 0.633; **4D**), or DMN-Salience network (T28) = 1.941, p = 0.0624; **Figure 4E**) for DOC patients with positive and negative responses to volitional tasks. However we did see significantly greater resting connectivity between FPCN and Salience networks (T(28) = 3.347, p = 0.0023, Bonferroni corrected p = 0.042; **Figure 4F**) in patients who responded to a volitional task compared to those who did not.

**Figure 4:**
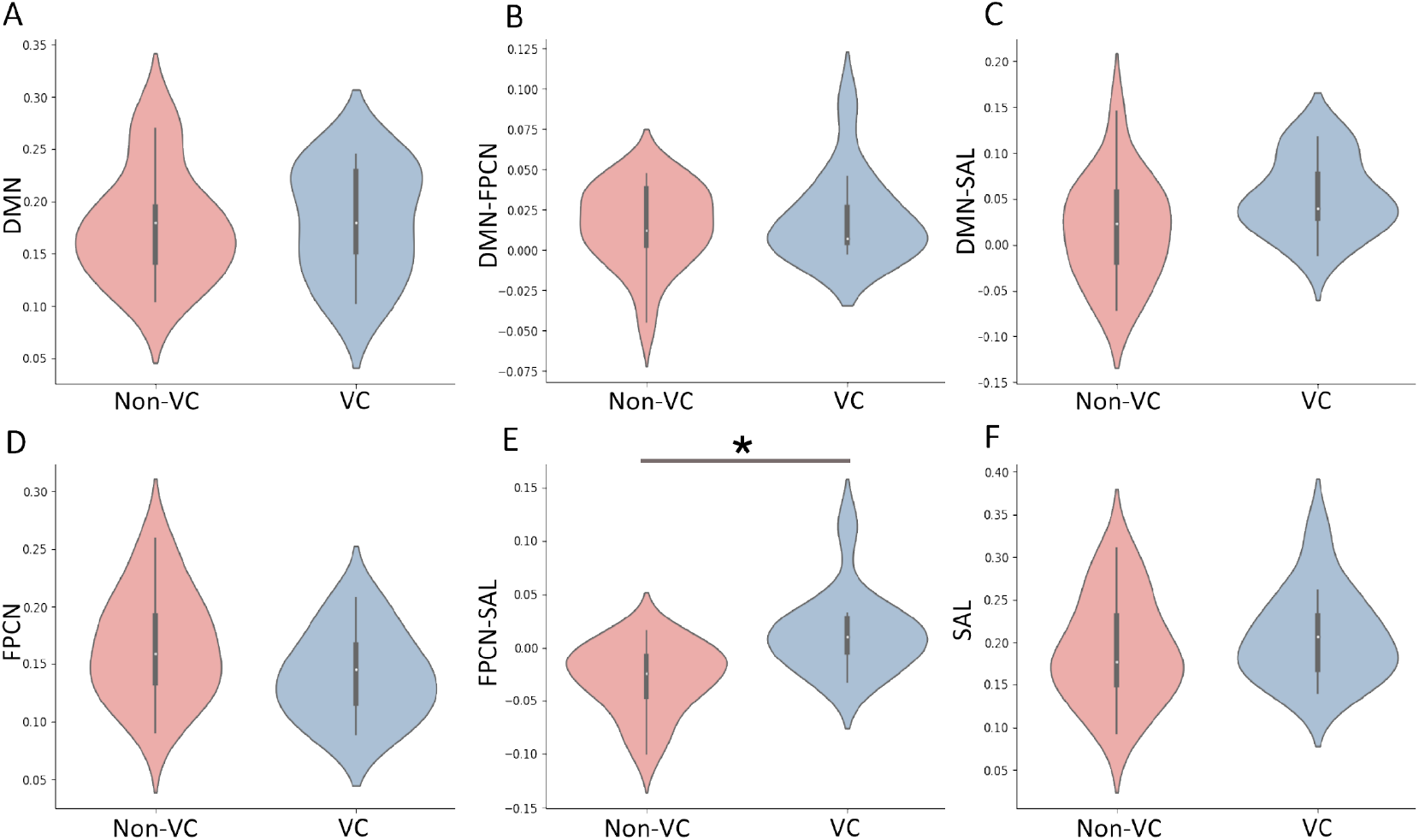
Violin plots of functional connectivity within and between networks in DOC with and without volitional capacity. **A** shows within DMN connectivity (T_(28)_ = 0.203, p = 0.84), **B** shows within FPCN connectivity (T_(28)_ = −1.681, p = 0.104), and **C** shows within Salience network connectivity (T_(28)_ = 0.732, p = 0.103) between positive and negative responders to volitional tasks. **D** shows between DMN-FPCN connectivity (T_(28)_ = 0.482, p = 0.633), **E** shows a between DMN-Salience network connectivity (T_28)_ = 1.941, p = 0.0624) and **F** shows a significant difference between FPCN-Salience network connectivity (T_(28)_ = 3.347, p = 0.0023, Bonferroni corrected p = 0.042) between positive and negative responders to volitional tasks.

### Classification Results

Next, we used a DGCNN to classify patients as VC or non-VC. We used each patient’s resting brain network connectivity graph as input to this algorithm. We also used individual brain networks (i.e. DMN, FPCN and Salience), as well as interactions between brain networks (i.e. DMN-FPCN, DMN-Salience, FPCN-Salience) as input to determine if a specific set of connections is more predictive of volitional capacity than another. We found that using all within and between network connections as input resulted in relatively high classification accuracy (test accuracy = 0.71; ROC-AUC = 0.79). However, when focusing specifically on within-FPCN connectivity as input, the accuracy of classifying VC patients substantially increased (test accuracy = 0.85; ROC-AUC = 0.92). Using within-network connectivity from DMN reduced accuracy, however, it was still well above chance (test accuracy = 0.62; ROC-AUC = 0.67). Within-Salience network connectivity showed very poor classification accuracy (test accuracy = 0.32; ROC-AUC = 0.29). For between-network connectivity, we found that using FPCN-Salience internetwork connections somewhat reliably classified patients (test accuracy = 0.71; ROC-AUC = 0.82). Both DMN-FPCN (test accuracy = 0.6; ROC-AUC = 0.66) and DMN-Salience (test accuracy = 0.55; ROC-AUC = 0.54) performed worse than FPCN-Salience connectivity. A complete list of results can be found in **Table 2**.

**Table 2:**
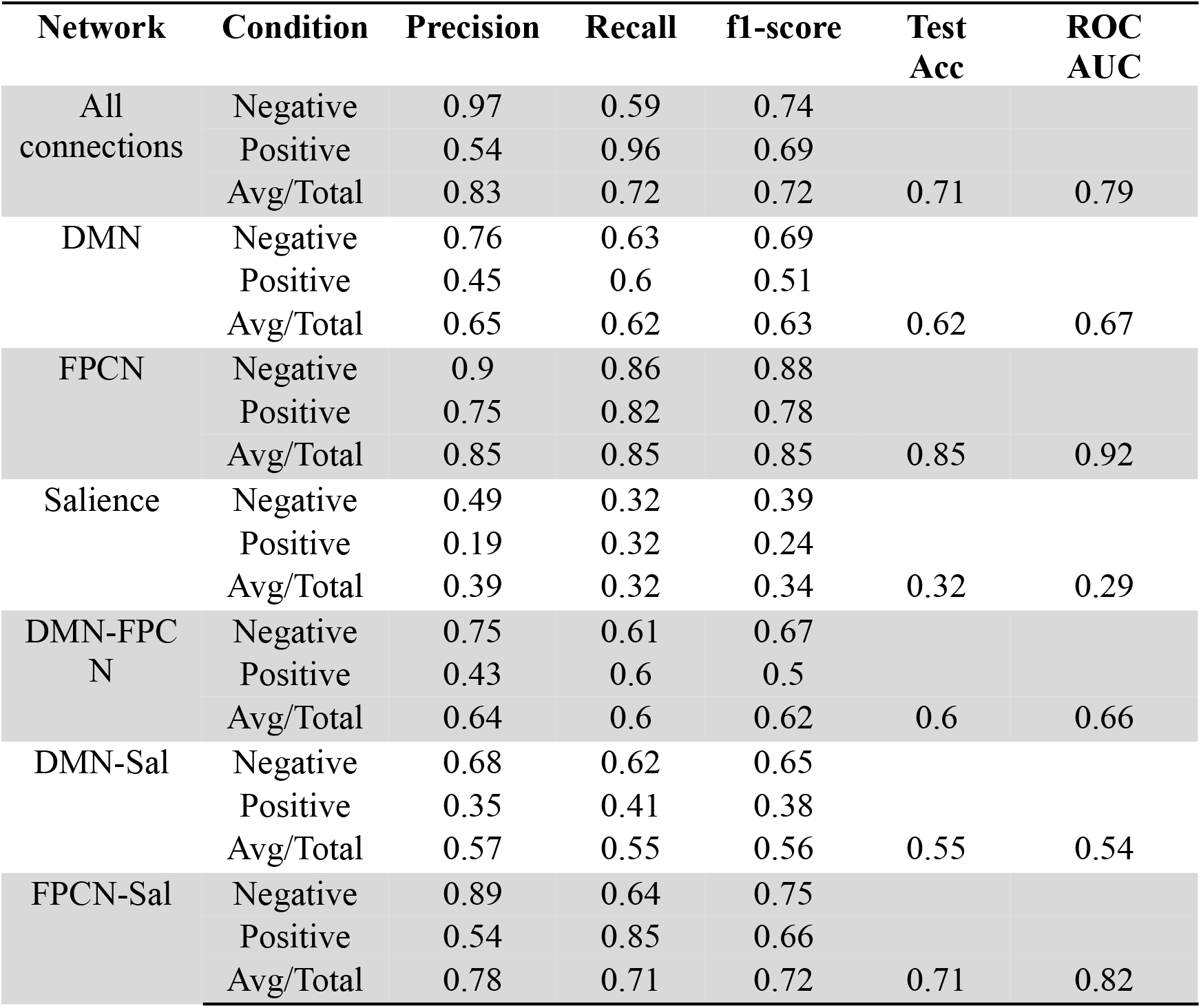
Classifying positive versus negative DOC responders to volitional tasks using a DGCNN on brain network data. Different brain networks were used as input to the DGCNN to determine which one contributed most to classification accuracy.

## DISCUSSION

The present study aimed to automate identification of volitional capacity in DOC patients using a resting-state functional connectivity-based deep learning classifier. This is the first study to use resting-state fMRI to predict volitional capacity in DOC patients using machine learning. It is well known that determining the diagnosis and prognosis of DOC patients is a challenging clinical problem. Understanding the degree of volitional capacity is vital to providing them with appropriate rehabilitation and can be greatly beneficial for their families. This makes the development of cutting-edge methodology critical to providing patients with the best care possible. To this end, our work builds upon the field by 1) using volitional capacity, instead of clinical diagnosis, as ground truth, and 2) by examining whether interactions within and between specific resting-state brain networks shown to be altered in DOC provide greater discriminatory power. This approach allowed us to not only build a useful statistical model that could be developed into a clinical tool but also provided a deeper understanding of the underlying network dynamics that predict each condition. As such, our model provides both useful clinical information regarding a patient’s volitional capacity beyond behavioural analysis and is neurobiologically interpretable.

We found that using only the FPCN as input to the DGCNN resulted in high classification accuracy (85% accuracy in held-out test data) as compared to other networks. We also found that connectivity between FPCN and Salience networks is significantly greater in patients capable of performing a volitional task. Patients in the VC group had significantly greater CRS-R scores than patients in the non-VC group. This is not surprising, given that patients with higher CRS-R scores are by definition more aware and wakeful, making it easier for them to respond to tasks. However, it is important to note that 6 of the 13 VC patients had CRS-R scores near or below the mean score for non-VC patients. This is critical, as this subset of patients may be incapable of physically responding to various motor commands from the CRS-R, making brain-based measures of volition vital during their assessment. This highlights the limitations of purely behavioural assessments and demonstrates that the added value of complementing behavioural assessments using brain imaging with automated analysis methods when diagnosing DOC patients.

Using several complementary approaches, we found that connectivity both within and between the FPCN was best at differentiating DOC patients with and without volitional ability. First, we found significantly greater connectivity between FPCN and Salience networks in patients capable of performing at least one mental imagery task compared to patients that did not. We also found that within FPCN connectivity and FPCN-Salience connectivity served as the best features for the DGCNN, respectively. This result reflects many previous studies suggesting that FPCN is vital for executive function and cognitive control (Demertzi et al., 2014; Dosenbach et al., 2007; Naci, Cusack, Anello, & Owen, 2014). Naci et al. (2014) examined whether individuals share common neural processes using a plot-driven movie-watching task. They showed that one behaviourally non-responsive patient displayed significantly similar patterns to controls in response to the task in frontoparietal regions. Our results show that preservation of awareness is reflected in FPCN connectivity not only during task performance but during resting-state as well. This work also suggests that intact frontoparietal connectivity is important for the preservation of conscious awareness and in an individual’s ability to wilfully control conscious experience. Previous work suggests that the DMN has significantly reduced connectivity at deeper levels of unconsciousness (Boly et al., 2009; Demertzi et al., 2014; Di Perri et al., 2016; Vanhaudenhuyse et al., 2010). Most recently, Di Perri et al. (2016) demonstrated that both within and between-DMN connectivity significantly correlated with the level of consciousness of DOC patients. Our analyses did not reveal any significant differences between VC and non-VC in either within or between-DMN connectivity, suggesting that DMN connectivity may reflect clinical diagnosis, but not volitional capacity.

It is well known that acquisition of data for patients with DOC is limited, and this is compounded by our selection criteria including the need for both task and resting-state fMRI to be collected for each specific patient. Deep learning models require large datasets for training and often require tuning to find the best set of hyperparameters to fit the data (Krizhevsky, Sutskever, & Hinton, 2012; Lecun et al., 2015). To overcome this limitation, we used a novel application of dynamic functional connectivity to augment the amount of data our deep learning model would be trained on. Instead of creating brain network graphs for a subject’s entire resting-state scan, we used a sliding window approach that generated multiple graphs per patient. Though having data collected from more patients would be preferable, our approach allowed us to properly train and tune our deep learning model, resulting in a high test accuracy for FPCN as an input feature. Translation of this work into a clinical tool will likely require both data augmentation methods and gathering data from additional patients.

In conclusion, the present study used resting-state functional connectivity and a deep learning algorithm to differentiate DOC patients who are capable or incapable of responding to a volitional task during fMRI scanning. Though volitional ability was significantly greater in the VC group, 6 of the 13 patients with volitional capacity had CRS-R scores near or below the average score for non-VC patients, demonstrating the utility for brain imaging methods to complement behavioural assessment during diagnosis. Furthermore, we found using FPCN as an input feature to the classifier provided the best accuracy when classifying patients into these conditions. We also found significantly greater connectivity between two task-positive brain networks, namely the FPCN and Salience Network in patients who could perform a volitional task. These results show that FPCN connectivity is particularly important in preserving and maintaining volitional ability and highlights the potential for use of deep learning in providing a clinical tool for neurologists in assessing patients with DOC.

## AUTHOR CONTRIBUTIONS

MMC was involved in literature search, figures, study design, data collection, data analysis, data interpretation and writing. IP was involved in study design, data analysis, data interpretation and writing. JA was involved in study design, data collection and data interpretation. PF was involved in study design, data collection and data interpretation. GW was involved in study design, data collection and data interpretation. JP was involved in study design, data collection and data interpretation. DKM was involved in study design, data collection, data interpretation and writing. EAS was involved in literature search, figures, study design, data collection, data analysis, data interpretation and writing.

## ACKNOWLEDGEMENTS

The authors thank the radiographers at the Wolfson Brain Imaging Centre for their assistance in data acquisition. We are grateful to the patients, their families and carers for their participation.

## COMPETING INTERESTS

The authors that they have no conflict of interest authors declare.

## FUNDING

This work was supported by grants from the UK Medical Research Council [U.1055.01.002.00001.01 to JDP; the James S. McDonnell Foundation to JDP; the National Institute for Health Research (NIHR, UK), Cambridge Biomedical Research Centre and NIHR Senior Investigator Awards to JDP and DKM; The Canadian Institute for Advanced Research (CIFAR) to DKM and EAS; the Stephen Erskine Fellowship (Queens ‘College, Cambridge) to EAS; the British Oxygen Professorship of the Royal College of Anaesthetists to DKM. MC was supported by the Cambridge International Trust and the Howard Sidney Sussex Research Studentship. IP was supported by Downing College, University of Cambridge through a Treherne Studentship;This work was also supported by the Evelyn Trust, Cambridge and the EoE CLAHRC fellowship to JA and the NIHR Brain Injury Healthcare Technology Co-operative based at Cambridge University Hospitals NHS Foundation Trust and University of Cambridge.

## ABBREVIATIONS

DOC: Disorders of Consciousness
FPCN: Frontoparietal Control Network
DMN: Default Mode Network
DGCNN: Deep Graph Convolutional Neural Network
VC: Volitional Capacity
MCS: Minimally Conscious State
UWS: Unresponsive Wakefulness Syndrome

## Notes

### Competing Interest Statement

The authors have declared no competing interest.

